# Matching drug transcriptional signatures to rare losses disrupting synaptic gene networks identifies known and novel candidate drugs for schizophrenia

**DOI:** 10.1101/268094

**Authors:** Daniele Merico, Xiao Wang, Ryan K. C. Yuen, Stephen W. Scherer, Anne S. Bassett

**Affiliations:** Deep Genomics Inc., Toronto; The Centre for Applied Genomics (TCAG), Genetics and Genome Biology, The Hospital for Sick Children, Toronto; Department of Molecular Genetics, University of Toronto, Toronto; Clinical Genetics Research Program and Campbell Family Mental Health Research Institute, Centre for Addiction and Mental Health (CAMH), Toronto; Toronto General Research Institute, University Health Network, Toronto; Department of Psychiatry, University of Toronto, Toronto; McLaughlin Centre, University of Toronto, Toronto

**Author notes:** Equal contribution (Daniele Merico, Xiao Wang).

## Abstract

Schizophrenia is a complex neuropsychiatric disorder. The etiology is not fully understood, but genetics plays an important role. Pathway analysis of genetic variants have suggested a central role for neuronal synaptic processes. Currently available antipsychotic medications successfully control positive symptoms (hallucinations and delusions) largely by inhibiting the dopamine D2 receptors; however, these drugs have more limited impact on negative symptoms (social withdrawal, flat affections, anhedonia) and cognitive deterioration. Drug development efforts have focused on a wide range of neurotransmitter systems and other agents, with conflicting or inconclusive results. New drug development paradigms are needed. A recent analysis, using common variant association results to match drugs based on their transcriptional perturbation signature, found drugs enriched in known antipsychotics plus novel candidates.

We followed a similar approach, but started our analysis from a synaptic gene network implicated by rare copy number loss variants. We found that a significant number of antipsychotics (p-value = 0.0002) and other psychoactive drugs (p-value = 0.0004) upregulate synaptic network genes. Based on global gene expression similarity, active drugs formed two main clusters: one with many known antipsychotics and antidepressants, the other with various drug categories including two nootropics. We specifically recommend further examination of nootropics with limited side effects (*meclofenoxate*, *piracetam* and *vinpocetine*) for combination therapy with antipsychotics to improve cognitive performance. Detailed experimental follow-up is required to further evaluate other candidate drugs lacking an official nervous system indication, although, for at least a few of these, psychoactive effects have been reported in the literature.

## INTRODUCTION

Schizophrenia is a complex neuropsychiatric disorder, with prevalence in the general population estimated at 1% [Kahn-NatRev-2015] and age of onset around 19-25 years old [Birnbaum-NatNsc-2017]. Clinical diagnosis is based on “operational criteria” that mainly rely on symptoms, without any objective diagnostic test [Kahn-NatRev-2015] [DSM-5] [Birnbaum-NatNsc-2017]. Schizophrenia symptoms are clustered in three major domains: positive (hallucinations and delusions), negative (social withdrawal, flat affections, anhedonia) and cognitive deficit [Liddle-BrJPsy-1987] [Kahn-NatRev-2015] [DSM-5]. Historically, there has been a greater diagnostic emphasis on positive symptoms, but cognitive deficits are particularly important and can be recognized in the earlier prodromal phase [Kahn-JAMAPsy-2013] [Forsyth-TrendsCognSci-2017]. Currently available antipsychotics inhibit the dopamine D2 receptors, although many antipsychotics are also able to modulate other neurotransmitter receptors [Creese-Science-1976] [Seeman-Nature-1976] [Miyamoto-MolPsy-2012] [Kahn-NatRev-2015]. Antipsychotics successfully control positive symptoms, but have limited impact on negative symptoms and cognitive deterioration [Miyamoto-MolPsy-2012] [Kahn-NatRev-2015]. *Typical* and *atypical* antipsychotics mainly differ in their side effects: extrapyramidal symptoms more for typicals, endocrine and metabolic side effects more for atypicals. Serotonin 5-HT2A receptor inhibition, lower affinity for and faster dissociation from for the dopamine D2 receptor only partially explain the atypical antipsychotics’ different activity profile [Miyamoto-MolPsy-2012] [Keshavan-ProgNeurob-2017]. *Clozapine*, one of the most potent antipsychotics, is typically prescribed to patients who fail to respond to other drugs; its potency cannot be accounted for only by its dopamine D2 receptor inhibition activity [Miyamoto-MolPsy-2012].

The etiology and pathophysiology of schizophrenia are complex, with many aspects presenting conflicting theories and still actively researched [Kahn-NatRev-2015] [Birnbaum-NatNsc-2017] [Forsyth-TrendsCognSci-2017]. Genetic variation is an important contributor to etiology, with heritability estimates up to 80%. Common variants providing the largest contribution, with a highly polygenic architecture [PGC:GWAS-Nature-2014] [Kahn-NatRev-2015] [Genovese-NatNsc-2016] [PGC:CNV-NatGen-2017] [Birnbaum-NatNsc-2017]. However, pathway analyses of common and rare loss-of-function variants have shown significant enrichment in neuronal synaptic components and synaptic plasticity [Fromer-Nature-2014] [Purcell-Nature-2014] [PGC:pathways-NatNsc-2015] [Genovese-NatNsc-2016] [PGC:CNV-NatGen-2017] [Forsyth-TrendsCognSci-2017].

Unlike other neurodevelopmental or neurodegenerative disorders, brain anatomical studies have not revealed gross pathological changes [Kahn-NatRev-2015] [Birnbaum-NatNsc-2017]. While there is no evidence for neuron loss, imaging studies have consistently revealed loss of brain volume [Kahn-NatRev-2015] [Birnbaum-NatNsc-2017] [Forsyth-TrendsCognSci-2017], with more detailed studies reporting decreased dendritic spine density [Forsyth-TrendsCognSci-2017] or increased post-natal synaptic pruning [Sekar-Nature-2016]. Beyond the well-established role of dopaminergic D2 receptors for positive symptoms, other neurotransmitter systems have also been examined [Kahn-NatRev-2015]. Glutamatergic N-methyl-D-aspartate receptor (NMDAR) antagonists (e.g. ketamine) induce psychosis-like states as well as cognitive deficits, suggesting an important role for the glutamatergic system [Miyamoto-MolPsy-2012] [Howes-JPsypharm-2015]. Accordingly, common and rare loss-of-function genetic variants have suggested glutamatergic dysfunction [PGC:GWAS-Nature-2014] [Fromer-Nature-2014] [PGC:CNV-NatGen-2017], while gene expression studies have suggested downregulation of glutamatergic synaptic components [Forsyth-TrendsCognSci-2017]. However, drug development results directly targeting the glutamatergic system have been mixed, and no pharmacological treatment modulating this system has been approved so far [Miyamoto-MolPsy-2012] [Kahn-NatRev-2015] [Howes-JPsypharm-2015] [Beck-Psychopharm-2016] [Keshavan-ProgNeurob-2017]. Abnormal GABAergic function has been consistently reported by multiple studies, but it is not clear if it merely reflects secondary adaptations and no approved schizophrenia drug directly targets GABAergic transmission either [Miyamoto-MolPsy-2012] [Kahn-NatRev-2015] [Birnbaum-NatNsc-2017]. Only combination of serotonergic modulation with antipsychotics has shown some promising results for negative symptoms [Miyamoto-MolPsy-2012] [Kahn-NatRev-2015] [Keshavan-ProgNeurob-2017]. Anti-inflammatory, antioxidants, neuroprotective agents and growth hormones have also been considered, but several have failed and none has yet completed clinical trials successfully [Miyamoto-MolPsy-2012] [Kahn-NatRev-2015] [Beck-Psychopharm-2016].

Lack of success can be attributed to the intrinsic complexity of the human brain and the consequent difficulties with mechanistically resolving the etiopathology, further complicated by differences of brain and behaviour between humans and model organisms [Hyman-Cerebrum-2013] [Kaiser-NatMed-2015] [Keshavan-ProgNeurob-2017]. Nonetheless, new research paradigms may disclose new promising avenues [Hyman-Cerebrum-2013] [Keshavan-ProgNeurob-2017]. Targets informed by genetics offer particularly good opportunities for development of new drugs [Nelson-NatGen-2015]. To capitalize on the bounty of genetic results, a recent analysis used common variant association statistics to infer the genetically-imparted differential expression associated to schizophrenia risk; this was then matched to drug-induced transcriptional signatures [Lamb-Science-2006], finding an enrichment in known antipsychotics as well as novel candidate compounds [So-NatNsc-2017]. This repositioning strategy has previously yielded candidates demonstrating validity in animal models for non-neuropsychiatric disorders [Dudley-SciTransMed-2011] [Sirota-SciTransMed-2011]; it additionally offers the advantage of suggesting new drug-disease connection beyond known pharmacological mechanisms for drugs that have already been tested for safety in humans [Hodos-wiley-2016] [Keshavan-ProgNeurob-2017].

In this study, we followed a similar approach to So et al [So-NatNsc-2017]. However, we focused our analysis on the specific protein networks showing higher burden of rare copy number losses in schizophrenia patients compared to controls [PGC:CNV-NatGen-2017]. Variants causing loss-of-function by reducing dosage (such as rare copy number losses) are ideal for matching to transcriptional response signatures. Focusing on rare variants of large effects additionally ensures the presence of strong and clear biological effects. Moreover, adopting a network-informed approach enables to generalize beyond individual variants found in a few subjects while preserving biological specificity and thus addressing the problem of etiological heterogeneity in schizophrenia.

We first confirmed that pre-constructed sets of drugs expected to be related to schizophrenia (antipsychotics and psychoactives) are over-represented in drugs with transcriptional effects remediating synaptic network rare losses, suggesting the validity of this strategy. We then investigated overall similarity relations among drugs, discovering two clusters of particular interest: one enriched in antipsychotics and antidepressants, the other with different drug categories including two nootropics. Based on these results, we specifically suggest the evaluation of nootropics with limited side effects to improve cognitive deficit. We also evaluated candidates without nervous system indications, finding evidence of psychoactive effects in the literature for at least some of them.

## RESULTS

### Synaptic gene networks enriched in rare copy number losses in schizophrenia

The Psychiatric Genomics Consortium (PGC) schizophrenia cohort of 21,094 cases and 20,227 controls with rare (at ≤ 1% frequency) copy number variants (CNVs) was previously analyzed for gene-set burden. That analysis identified a Gene Ontology (GO) derived set of synaptic components and the curated activity-regulated cytoskeleton-associated complex (ARC) as the most enriched in rare losses. Using the union of these two sets, a physical protein-protein interaction (ppi) network was seeded from genes passing an inclusive false discovery rate (FDR) threshold for locus-specific burden [PGC:CNV-NatGen-2017]. In addition to the original full network (which will be referred to as *extended synaptic*), for this study we manually identified two nested subnetworks for the drug matching analysis: *core synaptic* and *core glutamatergic* (see Figure 1). This was motivated by two main observations. First, the gene network presents particularly dense interactions around *DLG1* (Discs large, drosophila, homolog of), including other postsynaptic scaffold organizers (*DLG2*, *DLGAP1*, *SHANK1*, *SHANK2*) and glutamatergic ionotropic receptors (*GRID1*, *GRID2*, *GRIN1*, *GRIA4*), all with excess losses in cases. In addition, the presynaptic adhesion gene *NRXN1* (Neurexin 1), which is connected to the postsynaptic core by its trans-synaptic interaction with neuroligins, was also added to core glutamatergic (together with its immediate interaction neighbours) because it is important for glutamatergic synaptogenesis and maturation [Graf-2004-Cell] [Südhof-Nature-2008]. Second, the more sparsely-connected network periphery at the opposite end of *DLG1* and *NRXN1* is populated mostly by genes that belong to recurrent and large CNV loci. Contribution to schizophrenia risk for specific genes in this group is more uncertain, in spite of their synaptic function and the robust association of the multigenic CNV loci they belong to. For example, synaptic genes *P2RX6*, *SNAP29* and *SEPT5* belong to the 22q11.21 deletion region. However, a network analysis of 22q11.21 deletion subjects with and without schizophrenia suggested copy number loss of *CLTCL1*, *P2RX6*, *RTN4R* and *SEPT5* as more likely implicated in schizophrenia risk [Bassett-AmJPsy-2017]. In addition, *SEPT5*, but not *P2RX6* and *SNAP29*, is predicted haploinsufficient based on genetic constraint [ExAC-2016-Nature] (ExAC pLI = 0.86, 5.8E-07 and 0.12 respectively, see also Figure 1). For this reason, genes in this peripheral region were excluded from the core synaptic network and retained only in the extended synaptic network.

**Figure 1.**
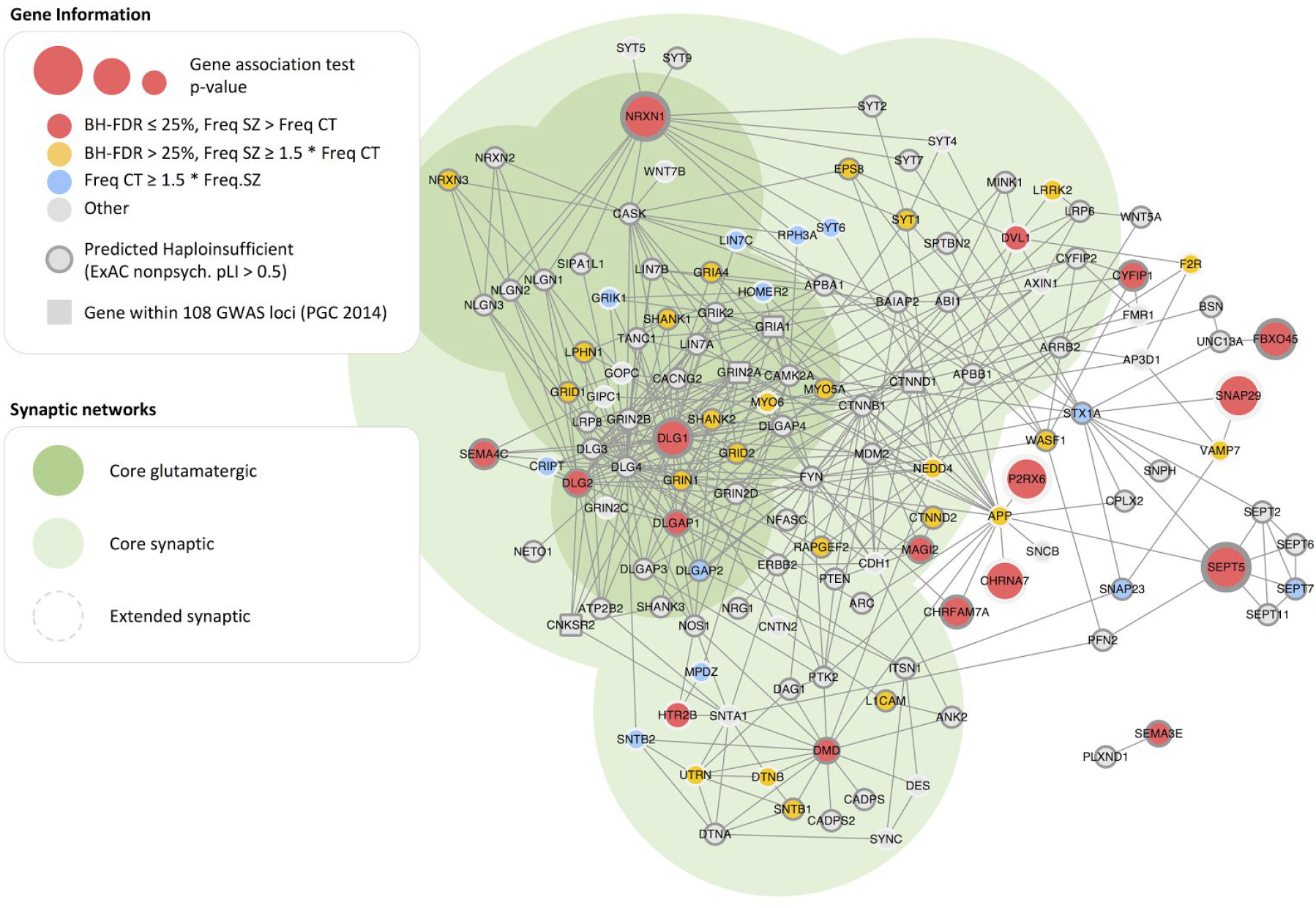
Nested synaptic networks: *core glutamatergic* (darker green shade), *core synaptic* (lighter green shade), *extended synaptic* (full network, white shade). Genes with excess losses in schizophrenia cases (Freq SZ ≥ Freq CT) are colored in red (locus FDR ≤ 25%) or gold (locus FDR > 25%, Freq SZ ≥ 1.5 Freq CT); genes with the opposite trend are colored in light blue. Predicted haploinsufficient genes (ExAC pLI, calculated after excluding psychiatric subjects) have a darker gray rim. Genes found within the 108 significantly associated GWAS loci are represented as squares instead of circles (*CNKSR2*, *CTNND1*, *GRIA1*, *GRIN2A*). See Supplementary Data Set 1 for detailed network gene annotations.

### Antipsychotics, psychoactive and nervous system drugs are over-represented in drugs increasing synaptic network expression

Similar to other studies [Ruderfer-LancetPsy-2016] [Gaspar-SciRep-2017] [So-NatNsc-2017], we used the Anatomical Therapeutic Chemical (ATC) classification system [ATC-URL] to a-priori define drug categories characterized by known indications relevant to schizophrenia and that had an available drug-induced differential expression signature in the Connectivity Map (CMap). This resulted in 28 antipsychotics (ATC ‘N05A: Antipsychotics’), 69 psychoactive drugs (union of ATC ‘N05: Psycholeptics’, which includes antipsychotics, anxiolytics, hypnotics and sedatives, and ‘N06: Psychoanaleptics’, which includes antidepressants, psychostimulants, nootropics and anti-dementia) and 142 nervous system drugs (ATC ‘N: Nervous System’). Of 1,217 drugs with CMap signatures, 803 drugs have an ATC classification.

CMap differentially expressed genes (DEGs) were generated from the original Affymetrix microarray image files (.CEL) using Robust Multi-array Average (RMA) [Irizarry-Biostatistics-2003] and Linear Models for Microarray Data (limma) [Ritchie-NAR-2015], with a single limma regression model capturing the drug effect on different cell types, at different doses and at different response times. Only HT Human Genome U133 Array data were processed, with probe-sets mapping to 13,814 unique genes. Requiring Benjamini-Hochberg False Discovery Rate (BH-FDR) ≤ 5%, the median number of up-regulated DEGs was 1,954 and the median number of down-regulated DEGs was 1,893; only 60 drugs had ≤ 100 DEGs and only 142 drugs had ≥ 6,500 DEGs (see Supplementary Data Set 2).

Using Gene Set Enrichment Analysis (GSEA) [GSEA-PNAS-2005], drugs were tested for enrichment in network gene upregulation, which corresponds to the opposite effect of copy number loss. For each network, we considered all network genes (*all*), or only network genes with excess losses in schizophrenia cases (*implicated*, coloured in red or gold in Figure 2). Each drug thus received a GSEA normalized enrichment score (NES), p-value and BH-FDR for each synaptic network and for each gene selection (all, implicated). Drugs with positive NES for a given set of network genes display an enrichment of upregulation in those genes compared to others, and thus are presumed to have a beneficial effect (see Supplementary Data Set 3 for detailed results).

**Figure 2.**
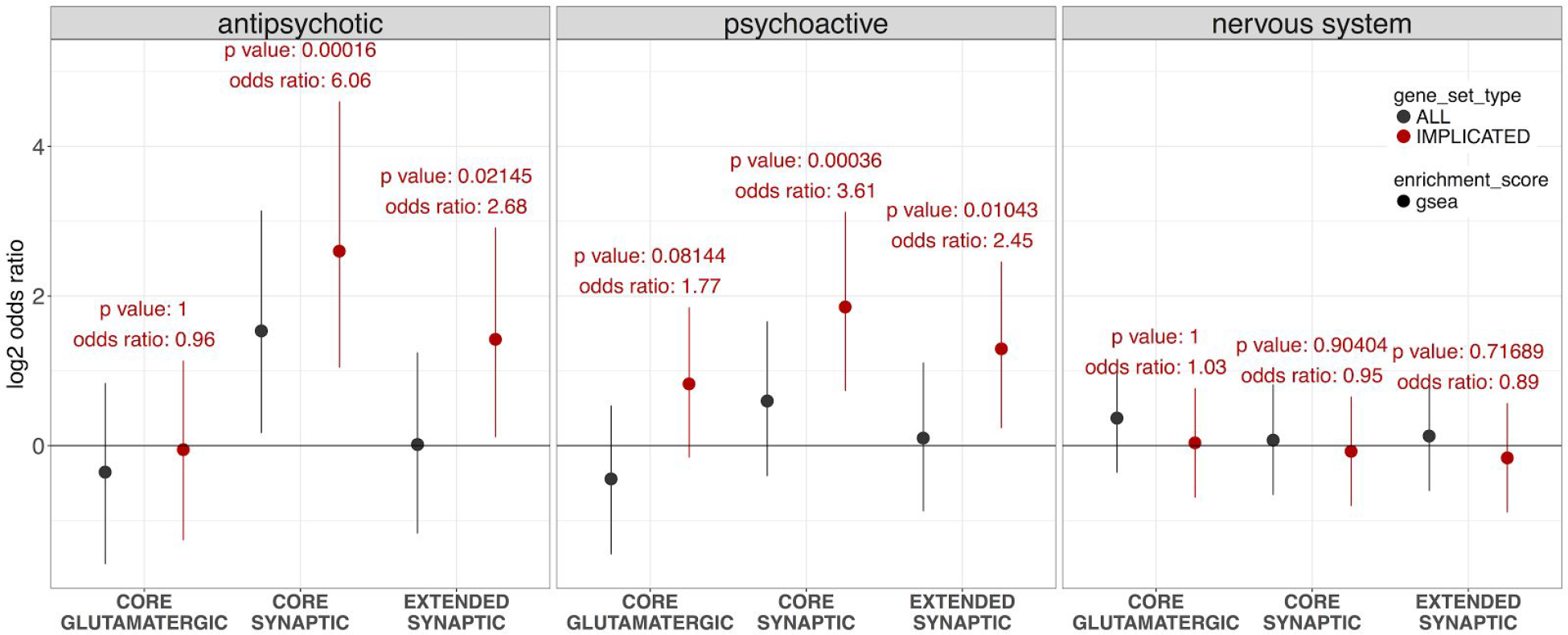
Over-representation in drugs with positive NES for the three pre-constructed drug categories (antipsychotics, other psychoactives and other nervous system), for each of the three nested synaptic networks (core glutamatergic, core synaptic, extended synaptic), when considering all network genes (*ALL*) or only genes with excess losses in schizophrenia subjects (*IMPLICATED*, corresponding to red and gold genes in Figure 1). Odds-ratio point estimates are depicted as filled circles and 95% confidence intervals are depicted as vertical lines; the y axis presents odds-ratios in log2 scale, whereas text labels present odds-ratios in linear scale.

Based on GSEA enrichment results, drug categories were then tested for over-representation of drugs with positive NES, using Fisher’s Exact Test. We tested psychoactive drugs removing antipsychotics and nervous system drugs removing psychoactive ones, to ensure that signal was not driven only by a small and specific subset of drugs within the larger one. Antipsychotics and other psychoactives displayed a robust over-representation in positive NES for the core synaptic network, and specifically for the implicated gene subset, with a non-significant trend towards a larger effect size in antipsychotics (antipsychotics, implicated: p-value = 0.00016, OR = 6.06; other psychoactives, implicated: p-value = 0.00036, OR = 3.61; see Figure 2 and Supplementary Table 1). Over-representation results were similar for the extended synaptic network, but with a non-significant trend towards smaller effect sizes than core synaptic. The core glutamatergic network (implicated subset) displayed borderline significant over-representation for other psychoactives (p-value = 0.08144, OR = 1.77), but not for antipsychotics (p-value = 1, OR = 0.96). Other nervous system drugs did not display any significant over-representation (p-value > 0.1).

Similar results were obtained when using a Wilcoxon rank-sum enrichment test instead of GSEA (see Supplementary Figure 1, Supplementary Table 2 and Supplementary Data Set 3). Setting more stringent enrichment thresholds (e.g. GSEA NES > 0 and BH-FDR < 25%) resulted in very large effect size confidence intervals and no significant over-representation, although effect size point estimates followed a trend similar to the main analysis (data not shown). Since over-representation results for the implicated gene subset were always better, we used this subset for all subsequent analyses.

We further investigated drug category over-representation following a more unbiased approach. When considering ATC first-level categories (14, corresponding to organ systems), only core synaptic presented significant over-representation (FDR ≤ 10%), and only for nervous system drugs. When considering ATC second-level categories (93), only core synaptic presented significant over-representation, for psycholeptics and psychoanaleptics. When considering ATC third-level categories (267), again only core synaptic presented significant over-representation, for antipsychotics. To maximize power, we also tested only the subset of third-level categories that are nervous system drugs. Core synaptic was found significantly over-represented in antipsychotics, antidepressants (N06A, 28 drugs) and hypnotics and sedatives (N05C, 7 drugs), whereas core glutamatergic and extended synaptic did not present any significant over-representation at FDR ≤ 10%. Antiepileptics (N03A,10 drugs) and nootropics (N06B, 4 drugs) ranked in the top 5 for all three networks, but had borderline or insignificant nominal p-values and fairly high BH-FDR (see Supplementary Data Set 4).

Antipsychotics and psychoactive drugs presented a similar number of drug-treated samples and significantly up-regulated genes (BH-FDR ≤ 5%) compared to other drugs, suggesting over-representation is robust to variations in experimental design (see Supplementary Table 3). These results validated our choice of the pre-constructed drug categories relevant to schizophrenia, with a striking suggestion that most antipsychotics (at least partially) revert the effect of rare loss copy number variants on the synaptic networks.

### Global similarity relations among drugs and prioritized drugs with nervous system activity

Drug similarity based on differential gene expression was quantified using Spearman correlation, followed by t-distributed stochastic neighbor embedding (t-SNE) projection to two dimensions for visualization [vanderMaaten-JMLR-2008]. When two drugs are in close proximity in the t-SNE plot, their differential profiles are expected to have a large and positive correlation. We manually identified two drug clusters over-represented in positive NES values for the core synaptic and core glutamatergic network (102 and 109 drugs respectively, see Figure 3a-b). Similar results were obtained using a Jaccard similarity for DEGs with BH-FDR ≤ 5%, suggesting this analysis is robust to the choice of the similarity metric (see Supplementary Figure 2). We tested the two clusters for over-representation in first-, second-and third-level ATC categories. We found the first cluster strongly over-represented in antipsychotics (15/28, OR = 14.54, BH-FDR ≤ 1.5 10^−5^%) and antidepressants (9/28, OR = 5.57, BH-FDR ≤ 2.4%), but also in non-nervous system categories such as gynecological anti-infectives and antiseptics (ATC G01, 8/24, OR = 5.83, BH-FDR ≤ 2.4%), antifungals for dermatological use (ATC D01, 6/17, OR = 6.25, BH-FDR ≤ 7.2%) and non-selective calcium channel blockers for cardiovascular use (ATC C08E, 3/4, OR = 33.51, BH-FDR ≤ 8.1%). Interestingly, there was also a borderline significant under-representation in anti-infectives for *systemic* use (ATC J, OR = 0.31, BH-FDR ≤ 37%), specular to the over-representation in anti-infectives for *topic* use. This cluster was therefore labelled *APAD* (AntiPsychotics and AntiDepressants). The second cluster was not significantly over-represented in any category (at BH-FDR ≤ 10%), even when using different cluster boundaries. The most nominally significant nervous system sub-category was psychostimulants and nootropics (ATC N06B, 2/4, OR = 8.52, p-value = 0.0580), and thus the cluster was labelled *MXNT* (MiXed drugs and NooTropics) (see Supplementary Data Set 5). Both clusters are over-represented in drugs with positive NES for the core glutamatergic and core synaptic network (APAD cluster: core glutamatergic p-value = 0.00258, OR = 1.89, core synaptic p-value = 4.31e-10, OR = 4.19; MXNT cluster: core glutamatergic p-value = 1.30e-11, OR = 3.85, core synaptic p-value = 2.38e-08, OR = 3.05). However, the APAD cluster, but not the MXNT cluster, displayed significantly smaller NES values for the glutamatergic network compared to core synaptic (APAD p-value = 8.06 10^−07^, MXNT p-value = 0.4880, Wilcoxon rank-sum test). Given the potential complementarity between antipsychotics and nervous system drugs inducing specific upregulation of the core glutamatergic network, we investigated various nervous system subcategories with some degree of over-representation in positive NES for core glutamatergic: none had a degree of clustering comparable to antipsychotics, antidepressants and nootropics (see Supplementary Figure 3).

**Figure 3.**
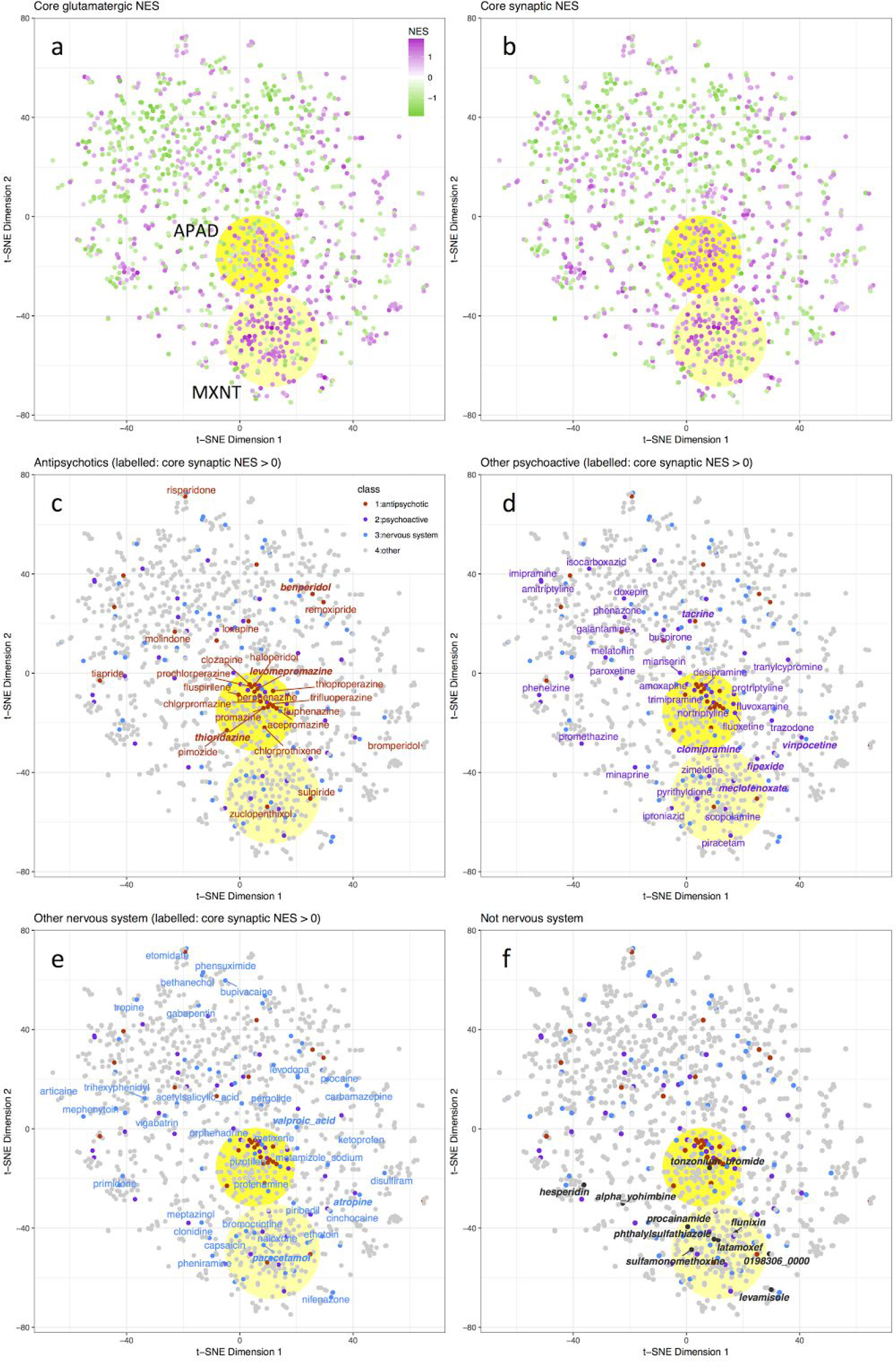
Global similarity relations among drugs based on differential expression Spearman correlation and t-SNE dimensionality reduction. All plots represent the same drugs with the same coordinates, but graphically displaying different attributes and drug subsets. In panels (a-b) drugs are coloured based on core glutamatergic (a) or core synaptic (b) NES: magenta for positive NES (enrichment in upregulation), green for negative NES (enrichment in downregulation). In panels (c-f) drugs are coloured based on drug categories: antipsychotics in red, other psychoactives in purple, other nervous system in blue, other drugs in gray or black. In panels (c-e), drug labels are displayed when core synaptic NES > 0, in bold italic for the subset in which NES and log2FC are positive for all three networks and core synaptic BH-FDR < 50% (c: antipsychotics; d: other psychoactives; e: other nervous system drugs). In panel (f), because of the large number of drugs without nervous system indication, labels are displayed only when NES and log2FC are positive for all three networks and core synaptic BH-FDR < 33%. In all panels, the APAD drug cluster (over-represented in antipsychotics and antidepressants) is represented as a full yellow shade, whereas the MXNT cluster (miscellaneous drugs with borderline over-representation of nootropics) is represented as a lighter yellow shade.

The vast majority of typical antipsychotics categorized as *phenothiazines* are part of the APAD cluster (10/12, see Table 1), suggesting a strong correlation between chemical structure and differential expression response. The majority of phenothiazines also have positive core synaptic NES, but not at a higher rate than other antipsychotic chemical classes (8/10 phenothiazines, 14/16 other antipsychotics, Fisher’s Exact Test p-value = 0.62542; see Table 1). The cluster also includes the powerful atypical *clozapine*, whereas the other two atypical antipsychotics, *risperidone* and *remoxipride*, map to different places (see Figure 3c). Other commonly used atypical antipsychotics (*aripiprazole*, *olanzapine*, *quetiapine*) are not represented in the CMap data-set. The antipsychotics *benperidol* (butyrophenone), *levomepromazine* and *thioridazine* (phenothiazines) have a positive effect on all three synaptic networks and pass a more stringent threshold for the core synaptic (positive NES and log2FC sum in all three networks and core synaptic GSEA BH-FDR < 50%, see Table 1 and Table 2). It is worth noting that the *levomepromazine* has a reputation of being a “dirty drug” because it interacts with a variety of targets and the branded version of *thioridazine* was withdrawn because it caused cardiac arrhythmias [Purhonen-PharmDrugSaf-2012]. While clozapine does not pass these thresholds, it is still remarkable that it has positive NES and log2FC sum for both core synaptic and core glutamatergic (see Supplementary Data Set 3).

**Table 1.**
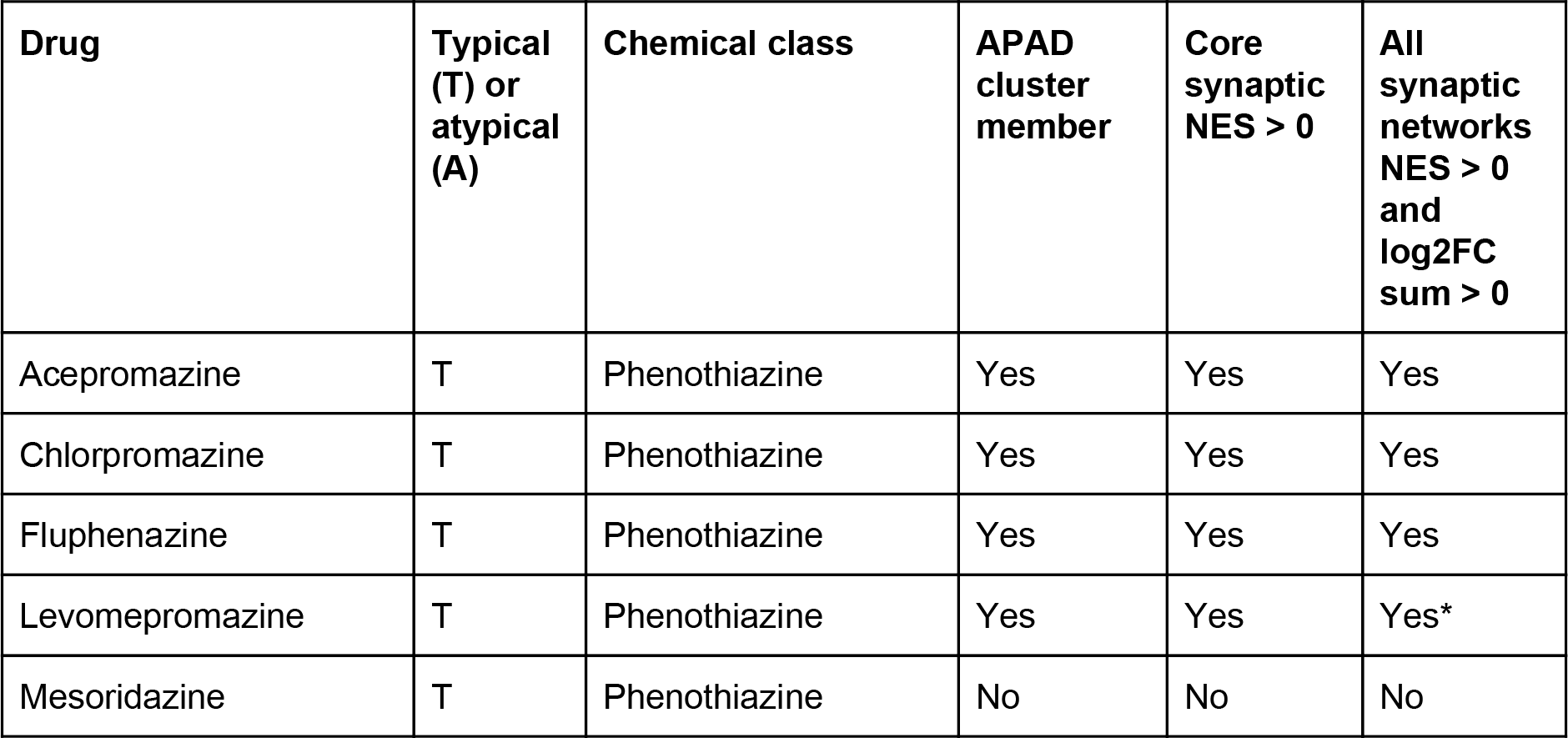
Antipsychotic drugs: typical/atypical classification, chemical structure type, t-SNE APAD cluster membership, synaptic network enrichment statistics. Drugs are sorted by chemical class and then alphabetically. In the ‘All synaptic networks NES > 0 and log2FC sum > 0′ column, ‘yes*’ indicates that the drug additionally has GSEA BH-FDR < 50% for the core synaptic network (these drugs have a bold italic font in Figure 3d).

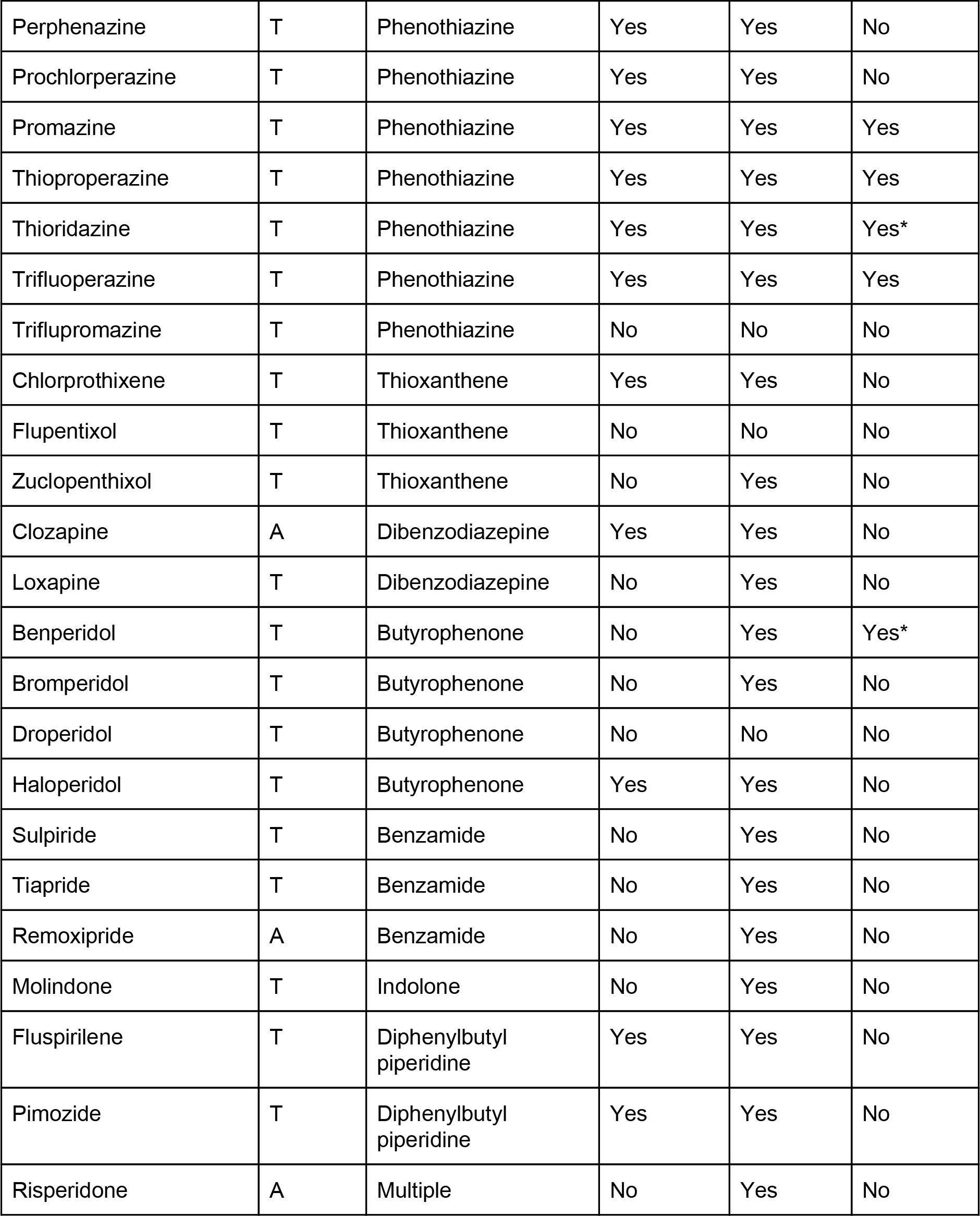

**Table 2.**
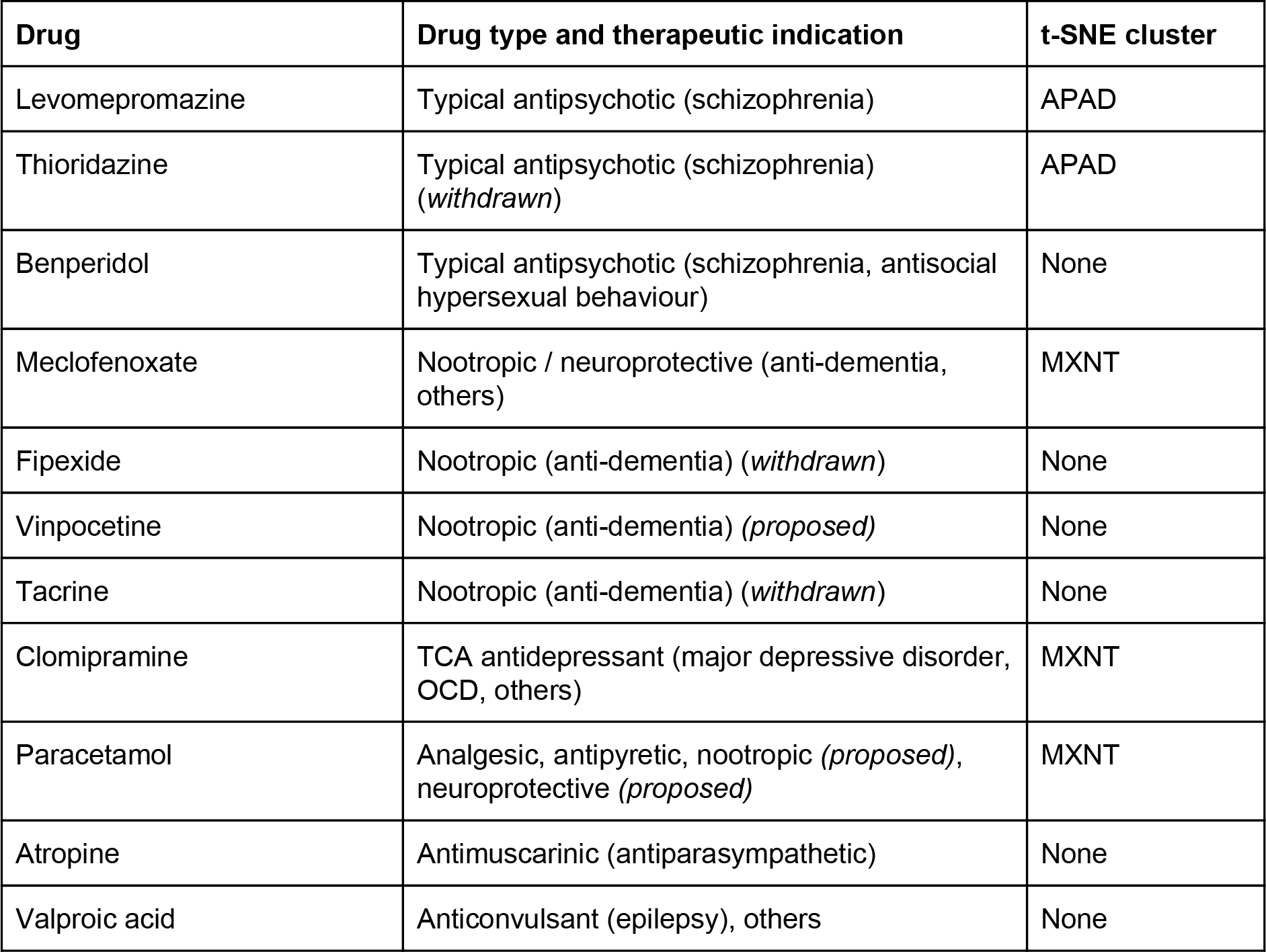
Prioritized nervous system drugs: these drugs have positive NES and sum log2FC for all synaptic networks, and also core synaptic network GSEA BH-FDR < 50%. The label ‘proposed’ is used whenever a therapeutic indication has been proposed only by preliminary research works but is not well established. Drugs are sorted following the order in the main text (first by indication, then by cluster membership, then alphabetically).

The APAD cluster also includes several antidepressants with positive core synaptic NES: the tricyclic antidepressants (TCA) *amoxapine*, *desipramine*, *protriptyline*, *nortriptyline* and *trimipramine*, and the specific serotonin reuptake inhibitors (SSRI) *fluoxetine* (*prozac*) and *fluvoxamine* (see Figure 3d). These drugs have in common the capability to inhibit serotonin reuptake, although TCAs also inhibit the reuptake of other neurotransmitters [Iversen-BrJPharm-2006]. TCAs have some degree of structural similarity to phenothiazines (the presence of two benzene rings), from which they were originally derived.

The MXNT cluster contains two nootropics, *meclofenoxate* and *piracetam*, both with positive core synaptic and core glutamatergic NES (see Figure 3d, Supplementary Data Set 3). *Meclofenoxate* is a cholinergic nootropic and neuroprotective agent reported to have beneficial effects for aging and other types of brain pathology [Marcer-AgeAgeing-1977] [Vaglenova-EurNPsyPharm-2001] [Nehru-BrainRes-2008] [Wang-PlosOne-2016]. *Piracetam* was shown to be effective to ameliorate ischemia-induced memory loss and cognitive deterioration; it also showed initial promising results in combination with *risperidone* to treat autism, whereas it was not effective for mild cognitive impairment and Down’s syndrome [Malykh-Drugs-2010]. An earlier report also suggested its use in schizophrenia [Noorbala-JClinPharmTher-1999]. The pharmacology of *piracetam* is complex and it has not been fully resolved: among various effects, it may act as an enhancer of glutamatergic and cholinergic transmission, but as an inhibitor of serotonergic and dopaminergic transmission [Malykh-Drugs-2010].

Considering the other two nootropics outside the MXNT cluster, *fipexide* and *vinpocetine*, it is interesting to note that both have positive effects on all three synaptic networks and core synaptic BH-FDR < 50% (see Table 2). In addition, while they are outside the MXNT cluster, they are in relative proximity (see Figure 3d) and they cluster more closely when using the Jaccard similarity (see Supplementary Figure 2d). *Fipexide* is a nootropic that was used for senile patients but later discontinued due to side effects such as liver necrosis [Bompani-CurrMedResOp-1986] [Durand-JHep-1992]. *Vinpocetine* use is not currently approved for clinical use, but preliminary findings relative to use for cognitive impairment and dementia are somewhat promising, with relatively limited side effects [Szatmari-Cochrane-2003]. Considering other MXNT cluster nervous system drugs, *clomipramine* and *paracetamol* have a positive effect on all three synaptic networks and core synaptic BH-FDR < 50% (see Table 2). *Clomipramine* is a TCA additionally used to treat obsessive-compulsive disorder (OCD); it has a mild dopamine D1, D2 and D3 receptor antagonist activity, which is particularly interesting considering its proximity to the APAD cluster (see Figure 3d) [Austin-BiolPsy-1991]. *Paracetamol* (also known as *acetaminophen*) is a commonly used over-the-counter analgesic and antipyretic, although its pharmacology is quite complex [Graham-Inflpath-2013]; a recent paper suggested a nootropic effect in humans, potentially related to interaction with serotonergic and cannabinoid systems, and supported by a previous animal study [Ishida-JPsychopharm-2007] [Pickering-DrugDesDevelTher-2016]. *Paracetamol* can also act as neuroprotective agent thanks to its antioxidant effect [Tripathy-JNeuroInfl-2009].

Considering nervous system drugs outside of the two clusters, *tacrine* is specifically worth a mention. *Tacrine* is a cholinestarase inhibitor used as nootropic in dementia (although it is classified only as anti-dementia in the ATC system); a relatively recent meta-analysis showed lack of significant improvements for Alzheimer Disease patients [Raina-AnnIntMed-2008] and tacrine was later withdrawn because of hepatotoxic side effects [Deardorff-DrugsAging-2015].

### Prioritized drugs without known nervous system activity

To identify top candidates lacking known nervous system activity, we set slightly more stringent criteria than for nervous system drugs: we required positive NES and sum log2FC in all synaptic networks plus core synaptic GSEA BH-FDR ≤ 33%. This resulted in 10 drugs, of which five are part of the MXNT cluster, two are not part but are in close proximity of MXNT cluster, and only one is part of the APAD cluster (see Table 3 for details). *Hesperidin* is particularly interesting, as it was also reported in the schizophrenia GWAS drug repositioning analysis [So-NatNsc-2017].

**Table 3.**
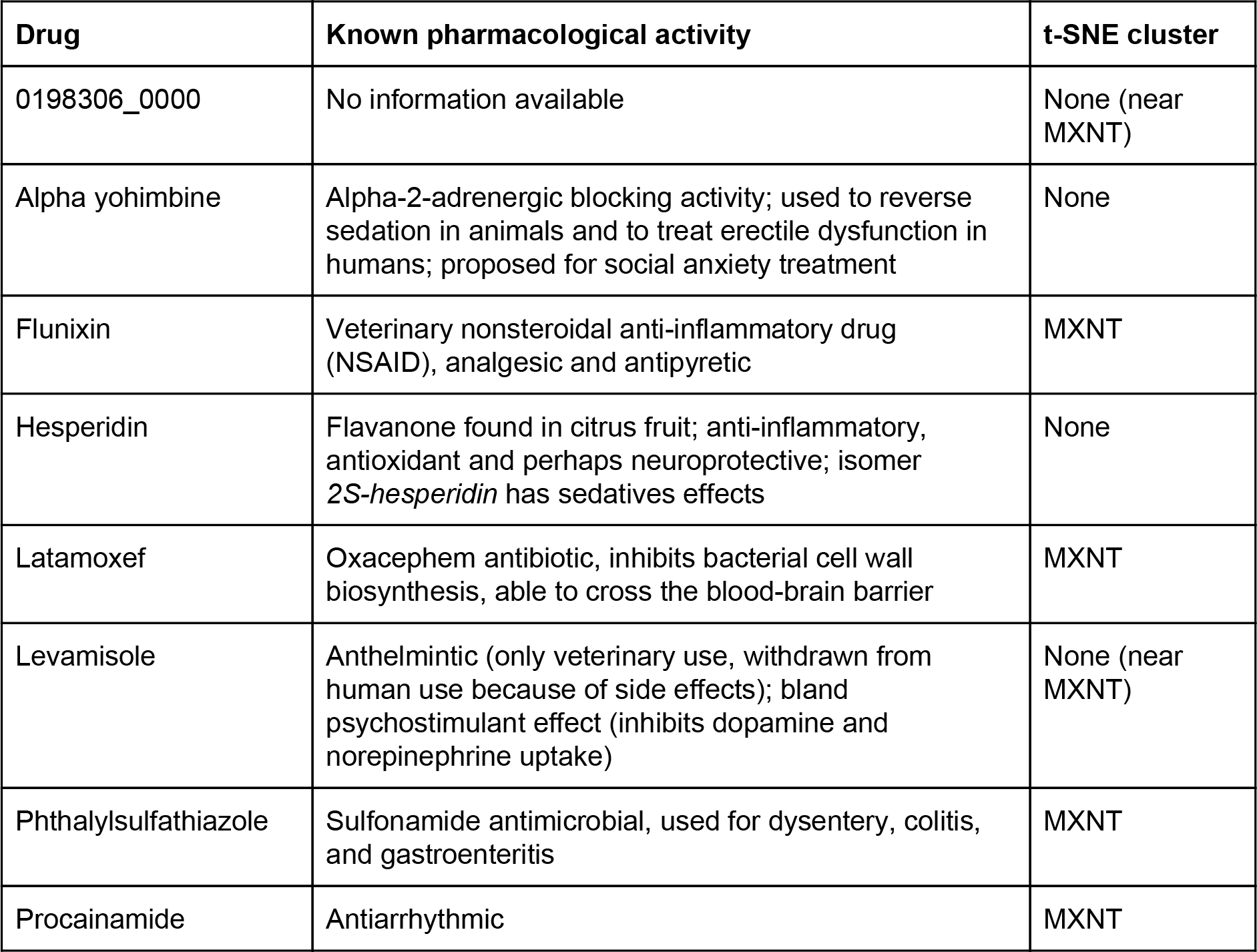
Top candidate drugs without nervous system activity, which have positive NES and sum log2FC for all synaptic networks and ‘core synaptic’ GSEA BH-FDR ≤ 33%.

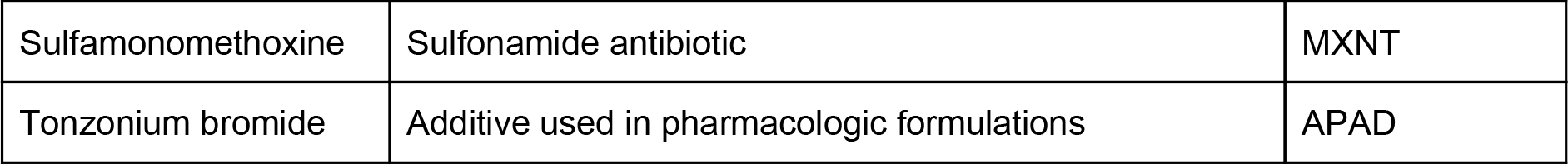

*Hesperidin* is a flavanone glycoside found in citrus fruit, with anti-inflammatory, antioxidant and perhaps neuroprotective properties [Garg-PhytRes-2001] [Parhiz-PhytRes-2015]. In addition, like other naturally occurring flavonoids, its isomer *2S-hesperidin* has sedatives effects [Fernández-EurJPharm-2006]. It co-localizes with *primidone* in a small cluster with positive core synaptic and core glutamatergic NES (see Figure 2f); *primidone* is an anticonvulsant barbiturate used in the past for seizures, which was also experimentally used for controlling aggression in psychotic patients [Monroe-JNervMentDis-1975]. Of other drugs, only *alpha yohimbine* and *levamisole* have known effects that could be of some interest to schizophrenia pharmacology (see Table 3). *Yohimbine* is a natural occurring indole alkaloid acting as an alpha adrenergic antagonist. It is used to reverse sedation in veterinary medicine [Zeiler-EqVetEd-2015] and to treat erectile dysfunction in humans [Andersson-PharmRev-2001]; it has also been recently proposed for treatment social anxiety disorder [Smits-BiolPsy-2014]. *Levamisole* is an anthelmintic that exerts its action by stimulating ionotropic acetylcholine receptors and causing worm paralysis; it is currently used in veterinary medicine, but it was withdrawn from human use because of its side effects. It also exhibits a bland psychostimulant activity by inhibiting dopamine and norepinephrine reuptake, and its metabolite aminorex has a more pronounced psychostimulant effect [Hofmaier-NeurochemInt-2014]. *Levamisole* clusters with the analgesic *nifenazone*, but the two drugs do not appear to have similar pharmacological targets.

### DISCUSSION

Dopaminergic D2 receptor inhibition successfully controls positive symptoms, but has more limited effect for cognitive deficit and negative symptoms, which are also critical for patient’s recovery and quality of life [Kahn-NatRev-2015]. After the successful adoption of typical and atypical antipsychotics in the 1950s-1970s, drug development efforts have focused on a wide range of neurotransmitter systems (dopaminergic, glutamatergic, serotonergic and cholinergic), but results have been generally inconclusive or negative and none has yet materialized into new treatments. A few other neuroligands and neuroprotective agents have also been investigated, without much success either [Miyamoto-MolPsy-2012] [Hyman-Cerebrum-2013] [Kahn-NatRev-2015] [Howes-JPsypharm-2015] [Beck-Psychopharm-2016] [Forsyth-TrendsCognSci-2017] [Keshavan-ProgNeurob-2017]. It is clear that new paradigms are needed for drug development. Genetic variation has a large contribution to schizophrenia etiology, and an increasing number of genetic loci have been associated to schizophrenia. Genetic findings may be particularly powerful for candidate drug generation. Recent studies have exploited the large schizophrenia cohort assembled by the PGC, mainly focusing on common variant genome-wide association (GWAS) results and experimentally-measured drug-protein interactions [Lencz-MolPsy-2015] [DeJong-JPsychopharm-2016] [Ruderfer-LancetPsy-2016] [Gaspar-SciRep-2017]. Reassuringly, several of these studies have found over-representation of antipsychotics targets among top-associated genes [DeJong-JPsychopharm-2016] [Ruderfer-LancetPsy-2016] [Gaspar-SciRep-2017]. One of these studies also investigated very rare and disruptive variants from whole exome sequencing, finding convergent results with GWAS loci and suggesting these rare variants may be also implicated in antipsychotic resistance [Ruderfer-LancetPsy-2016]. However valuable, neither GWAS associations nor physical drug-protein interactions provide information on effect directionality. Reliance on physical drug-protein interactions can additionally limit results to already known and well-characterized pharmacological properties. To overcome these issues, a recent study used common variant association results to infer differential expression associated to schizophrenia risk; drugs were then prioritized for their capability to produce inverse differential expression patterns in cell line assays, thus modelling directionality at the drug and genetic effect level. Candidates found using this strategy were significantly enriched in antipsychotics, but also included other compounds that will need to be further investigated experimentally [So-NatNsc-2017].

We followed a similar approach, but started our analysis from a synaptic gene network implicated by rare copy number loss variants in schizophrenia patients compared to controls.

Rare variants have larger effect sizes (significantly associated rare losses typically had case-control odds ratio ≥ 10): this helps establishing clear causal relations between genetic variation and biological consequences, which is more challenging for common variants because of linkage disequilibrium and small effect sizes [Harrison-JPsypharm-2015]. Rare copy number losses also directly suggest consequent gene expression changes without requiring complicated inference models. On the other hand, in presence of multigenic copy number changes, it may not be clear which genes contribute to disease risk. To circumvent this problem, we capitalized on previously published network analysis results, pinpointing to a synaptic gene network: other rare and common variant pathway analyses have implicated similar sets of genes [Fromer-Nature-2014] [Purcell-Nature-2014] [PGC:pathways-NatNsc-2015] [Genovese-NatNsc-2016]. In addition to the full gene network (extended synaptic), we also considered two nested subnetworks (core synaptic and core glutamatergic) for our analysis. Similar to findings based on common variants [So-NatNsc-2017], we found antipsychotics to be over-represented in drugs leading to synaptic network up-regulation and thus reversing the rare loss effects. We found a similar trend for the other psychoactive drugs, including antidepressants, nootropics and anti-dementia drugs, but not for other nervous system drugs. This is an important initial proof-of-concept, further demonstrating the validity of using cell line transcriptional perturbation signatures for pharmacology of CNS conditions.

We then investigated global similarity relations between drugs: we think this is a critical step to differentiate therapeutic opportunities that mimic the effect of antipsychotics or that act differently and thus may have a synergistic effect. We identified two major clusters: (1) APAD, strongly over-represented in antipsychotics (and specifically phenothiazines), but also antidepressants and other drugs; (2) MXNT, without any specific over-representation, but including two nootropics and other nervous system drugs. While clustering patterns (at least partially) reflect the known pharmacology, they also suggest the existence of more complex mechanisms: for instance, 9/24 antipsychotics with positive effects on the core synaptic network are not member of the APAD cluster, suggesting they induce different global transcriptional effects. The over-representation in antipsychotics but also antidepressants for the APAD cluster is intriguing, since schizophrenia presents a high genetic correlation to bipolar disorder and depression [Bulik-Sullivan-NatGen-2015]. Antidepressants and specifically serotonergic modulators have been previously investigated for schizophrenia treatment; preliminary results have suggested some benefit for negative symptoms but no clinical trials have been concluded successfully yet, and thus no antidepressant is included in treatment guidelines [Miyamoto-MolPsy-2012] [Kahn-NatRev-2015] [Keshavan-ProgNeurob-2017].

Since known antipsychotics have limited impact on negative symptoms and cognitive deficits, it is interesting to speculate if this is reflected somehow in our analysis results. The APAD cluster presented weaker enrichments than the MXNT cluster relative to the core glutamatergic network, corresponding to key components of glutamatergic transmission (glutamate ionotropic receptors, synaptic organizers and trans-synaptic adhesion proteins), similar to the overall trend for antipsychotics. For this reason, nootropics and other MXNT cluster drugs may results in a synergistic effect when used in combination with standard-of-care antipsychotics. Drugs with limited side effects, such as *meclofenoxate, piracetam* and *vinpocetine*, may represent a particularly promising opportunity for clinical translation. In fact, glutamatergic stimulation using *racetam* class nootropics has already been proposed based on GWAS findings [Lencz-MolPsy-2015]. Considering the small number of nootropics profiled in this data-set, investigation of other MXNT cluster drugs for nootropic effects may be particularly useful.

As far as other top-ranking drugs without a nervous system indication, follow-up experiments are necessary to demonstrate that they can have a psychoactive effect. Positive results are not unlikely, at least for a subset of these candidates: for instance, the anthelmintic *levamisole* was previously found to be also a bland psychostimulant. Among these candidate drugs, the citrus fruit flavonoid *hesperidin* may be particularly interesting, since it was also found as the second top hit (among 18) by matching transcriptional signatures to common variants [So-NatNsc-2017]. *Hesperidin* is expected act as a neuroleptic and/or neuroprotective agent. Further investigation of non-selective calcium channel blockers (over-represented in the APAD cluster) may also be promising: calcium channels have been implicated in schizophrenia by rare and common variants, and their modulation has been previously suggested as therapeutically promising [Lencz-MolPsy-2015].

These results are primarily limited by the number of compounds tested (1,217). A new CMap data-set has been very recently released, profiling a reduced number of genes and a much larger number of drugs [Subramanian-Cell-2017].

Reliance on cell lines for transcriptional signatures is another important source of potential false negatives, although systems more representative of in vivo brain function (such as neurons differentiated from iPS cells or brain organoids) likely present major challenges for reproducibility (of cell type composition and differentiation stage) compared to stable cell lines. Transcriptional perturbation signatures are used as proxy of cell state: additionally measuring neuronal cell phenotypes (like nervous signal transmission and synaptic formation or pruning) would be highly beneficial, but would also require a major experimental effort to test a sufficiently large number of drugs.

A more fundamental challenge is posed by the heterogeneous etiology of schizophrenia and the reliance, as a nosological construct, on clinical consensus rather than more mechanistic and biologically-rooted criteria. Developing experimental models of its key pathological processes, whether using human brain organoids or model organisms, is probably going to be one of the major challenges on the way towards successful drug discovery [Keshavan-ProgNeurob-2017]. We argue that variants of large effect that are able to recapitulate the disease and its known pharmacology in experimental models, such as the rare copy number losses in the synaptic networks described in this work, constitute a particularly valuable resource towards this goal.

## METHODS

### CMap gene expression data

The expression signatures were downloaded from the cmap database: https://portals.broadinstitute.org/cmap/. The full data-set consists of 7056 CEL files corresponding to three different array platforms (807 for HG-U133A, 6029 for HT_HG-U133A and 220 for HT_HG-U133A_EA). To avoid between array normalization, this study focused on the drugs profiled on the HT_HG-U133A platform, comprising a total of 5,242 treated and 787 untreated samples tested on three different cell lines (MCF7, PC3, HL60). The CEL files were imported using the R/Bioconductor *affy* package v1.54.0 and probe-set level gene expression summaries, normalized between samples, were generated using the rma method [Irizarry-Biostatistics-2003] as implemented the affy package [Affy-Bioinformatics-2004].

### Differential gene expression analysis

For each drug, the following generalized linear model was fit using the R/Bioconductor package *limma* v3.32.10 [Ritchie-NAR-2015]:

*Expr = b1 * combinedFactor* + *b2 * treatment* + *b3 * combinedFactor * treatment* where combinedFactor corresponds to the combined values of the cell line used for screening, the drug dose and the response time after treatment. Differential expression p-values and log2 fold-change (FC) were calculated based on the b2 coefficient.

### GSEA and Wilcoxon enrichment analysis

First, differential gene expression results were processed to convert probe-set identifiers to gene identifiers (NCBI Entrez-Gene), using the mapping table from the R/Bioconductor package *hthgu133a.db* v3.2.3. In presence of multiple probe-sets mapping to to the same single entrez gene id, only the most most differentially expressed probe-set was retained. Then, for each drug, GSEA pre-ranked version 3.0 (using default weighting) was used to analyze synaptic network enrichment in upregulation, providing in input genes ranked by limma moderated t-statistic (equivalent to ranking by-log10 (p-value) * sign (log2 FC)) and 6 gene-sets corresponding to each combination of network (core glutamatergic, core synaptic, extended synaptic) and gene subset (all, implicated). GSEA enrichment p-values were corrected for multiple testing using BH-FDR for each combination separately, across all drugs. For the Wilcoxon enrichment analysis, the GSEA pre-ranked test was replaced by a two-sided Wilcoxon rank-sum test as implemented in the R function *wilcox.test()*; the GSEA NES was replaced by an enrichment score (ES) calculated as the difference of the average of the gene inverse ranks for the gene-set and the other genes (where an inverse rank for N values is N for the largest value and 1 for the smallest); multiple test correction by BH-FDR was performed following the same logic as for the GSEA enrichment analysis.

### Drug category over-representation

Drug category over-representation in drugs with positive NES drugs and in drugs within t-SNE clusters was performed using a two-sided Fisher’s Exact Test as implemented by the R function *fisher.test()*; given a network of interest, the contingency matrix was defined as: (a) number of drugs within category (or within cluster) and NES > 0, (b) number of drugs within category (or within cluster) and NES < 0, (a) number of drugs outside the category (or outside the cluster) and NES > 0, (b) number of drugs outside the category (or outside the cluster) and NES < 0.

### Multiple test correction using BH-FDR

BH-FDR multiple test correction was performed using the R function *p.adjust(method = “BH”)*.

### t-SNE dimensionality reduction

Differential expression data was first converted to a similarity matrix using the Spearman correlation between limma moderated t-statistic vectors, calculated using the R function *cor(method = “spearman”)*. The same analysis was repeated using as similarity the Jaccard index between sets of differentially expressed genes at BH-FDR < 5%, averaging for every drug the Jaccard index separately calculated for upregulated and downregulated genes.

The similarity matrix was used to derived the 2d projection using to the *tsne()* function in the *tsne* R/CRAN package v0.1-3. [vanderMaaten-JMLR-2008]

## SUPPLEMENTARY ITEMS LIST

### Supplementary figures

- Supplementary Figure 1: over-representation in drugs with positive NES for the three pre-constructed drug categories, based on Wilcoxon enrichment analysis instead of GSEA
- Supplementary Figure 2: t-SNE results based on Jaccard similarity (note that the MXNT cluster is displayed in light orange instead of light yellow)
- Supplementary Figure 3: t-SNE plots highlighting nervous system drug sub-categories with positive core glutamatergic NES

### Supplementary tables

- Supplementary Table 1: drug category over-representation statistics based on GSEA enrichment, as displayed in Fig 2
- Supplementary Table 2: drug category over-representation statistics based on Wilcoxon enrichment, as displayed in Suppl Fig 1 (Wilcoxon ES replaces GSEA NES)
- Supplementary Table 3: sample and DEG counts for antipsychotics and psychoactive drugs compared to other drugs

### Supplementary datasets

- Supplementary Data Set 1: synaptic network gene annotations
- Supplementary Data Set 2: experimental design summary and DEG count for each drug
- Supplementary Data Set 3: GSEA and Wilcoxon signed rank enrichment statistics
- Supplementary Data Set 4: over-representation statistics for all level-1, level-2 and level-3 ATC categories, implicated gene subset
- Supplementary Data Set 5: APAD and MXNT cluster drug category over-representation results

## OTHER SECTIONS

### Contributions

RKCY and DM originally conceived the study; DM, AB and RKCY planned the study; XW and DM performed the analysis; DM, ASB, RKCY, XW and SWS interpreted the results; XW refactored and finalized the code repository; DM, ASB, XW and RKCY wrote the paper; all authors contributed to revising the paper.

**Conflicts of interest**

